# Pangenomes of Human Gut Microbiota Uncover Links Between Genetic Diversity and Stress Response

**DOI:** 10.1101/2024.04.17.589959

**Authors:** Saar Shoer, Lee Reicher, Yitzhak Pilpel, Eran Segal

## Abstract

The human gut microbiota has been the subject of many studies, establishing its central role in host health and disease. Interplay between mutations, horizontal gene transfer, natural selection, and genetic drift, lead to genetic diversity within these species, resulting in different phenotypes and effects on the host. Pangenome represents the collective gene pool of different strains of the same species. Here, we created pangenomes for 728 human gut prokaryotic species, holding four times more genes than the highest quality individual genome, showcasing the genetic diversity inherent in the human gut population. We find these species have a core set of about a thousand genes that defines them, distinct even between closely related species, and an accessory set of genes that are unique to the different strains. Furthermore, we show a spectrum of microbial behavior, while some species exhibit a saturated or “closed” pangenome, suggesting a limited set of genetic capabilities, others maintain an “open” pangenome, indicating elevated adaptability through genetic diversity. We discover that high strain variability is associated with the capacity of species to undergo sporulation, whereas low strain variability is associated with carrying genes that facilitate antibiotic resistances, suggesting different evolutionary strategies for survival taken by these microbes. We further map the landscape of antibiotic resistance genes across the human gut population, and find 237 cases of extreme resistance, predominantly of Enterobacteriaceae species, even to last resort antibiotics kept for cases where traditional treatments have failed. Lastly, we associate microbial strain level differences with human age and sex, exemplifying how the presence of specific genes in *Akkermansia muciniphila* and *Phocaeicola vulgatus* relate to host characteristics. Overall, our research provides a comprehensive overview of the evolution, genetic complexity and functional potential of the human gut microbiota, emphasizing its significant implications for human health and disease. The pangenomes and the antibiotic resistances map constitute a valuable resource for further scientific research and therapeutic advancements.

## Introduction

The human gut microbiota has been the subject of many studies, establishing its central role in host health and disease ^1^. Advancements in culture-independent technologies have revealed the vast genetic diversity, functional capacity, and dynamic nature of the human microbiota. Microbiota dysbiosis, imbalances in the composition and function of these microbes, has been linked to a spectrum of diseases including inflammatory, metabolic, hepatic, cardiovascular and even neurological disorders ^1^.

Within this microbial community, “species” is defined as a group of individuals forming a coherent genomic cluster ^2^. A widely recognized prokaryotic species delineation standard is 95% average nucleotide identity (ANI) ^2^. Despite the genetic coherence, broad phenotypic variability can be seen among strains of the same species. This subject is particularly well studied in the context of pathogenicity, where both commensal and pathogenic strains of the same species can be found. The traditional concept of “strain”, rooted in culture-dependent analysis, can not be directly transferred to modern culture-independent approaches, and a widely accepted biologically meaningful definition of a strain remains elusive ^2^. In this work a strain would refer to an assembled genome from a particular human. “Sub species” defined as a genetically or phenotypically distinct cluster of strains, can shed light on the evolutionary adaptability of species to their environment and explain heterogeneous interactions with the host ^2^.

The genetic diversity within microbial species arises from a continuous interplay of mutations, horizontal gene transfer, natural selection, and genetic drift ^3^. Mutations (substitutions, insertions, deletions, and inversions) introduce genetic changes through DNA damage, or errors in the processes of replication, recombination and repair. Horizontal gene transfer (through mechanisms of transformation, transduction, and conjugation) allows the rapid acquisition and dissemination of genetic material, significantly altering the genomic structure. Horizontal gene transfer can occur both within species and between distantly related species. However, it is generally more common within species or closely related species due to the higher likelihood of successful integration and expression of transferred genes ^4-5^. The subsequent fate of these genetic variations is determined by natural selection, favoring traits that enhance fitness on one hand, and genetic drift, which randomly alters traits’ frequency on the other.

A pangenome is the collective set of genes observed across all strains of a given species. In this work, we created pangenomes for 728 human gut prokaryotic species, a resource describing the genetic diversity within this environment, that can improve reference-based analyses that are inherently limited by the content of individual strains’ genomes. Additionally, we mapped the landscape of antibiotic resistance genes within the human gut microbiota, as species that are commensal in this environment are often pathogenic elsewhere. We anticipate that such knowledge may help fight this rising global concern.

## Results

### 728 human gut prokaryotes pangenomes

In a previous study, we assembled prokaryote genomes from 51,052 human stool metagenomic samples and added genomes from other published sources, to form a set of 241,118 genomes (methods) ^6^. To ensure diversity and unbiased representation, we limited the dataset to one genome from the same human individual in each species, making the terms “genome” and “strain” interchangeable within our dataset. We clustered this set based on genomic distances between pairs, and chose the best genome for each species cluster based on quality parameters (methods). In this way, we constructed strain sets for 3,594 human gut prokaryotic species.

In the current study, we created pangenomes for 728 species that have 220 strains in their respective clusters, and often based on 70 strains (median, range 20-9,834, methods). The vast majority of these species are bacteria, with only three belonging to the archaea domain. Our typical pangenome holds 9,158 genes (median, range 1,432-98,890, with two outliers of 222,346 and 410,781), representing a four fold increase compared to the highest quality species genome within the cluster, typically holding 2,414 genes (median, range 864-6,977, Figure 1). If only 20 strains were used to construct the pangenome of each species, the typical genes fold change would be two (methods, Figure1 insert). This significant gene expansion highlights the high genetic diversity inherent in the bacterial population of the human gut, and the pangenomes provide means to research it.

**Figure 1.**
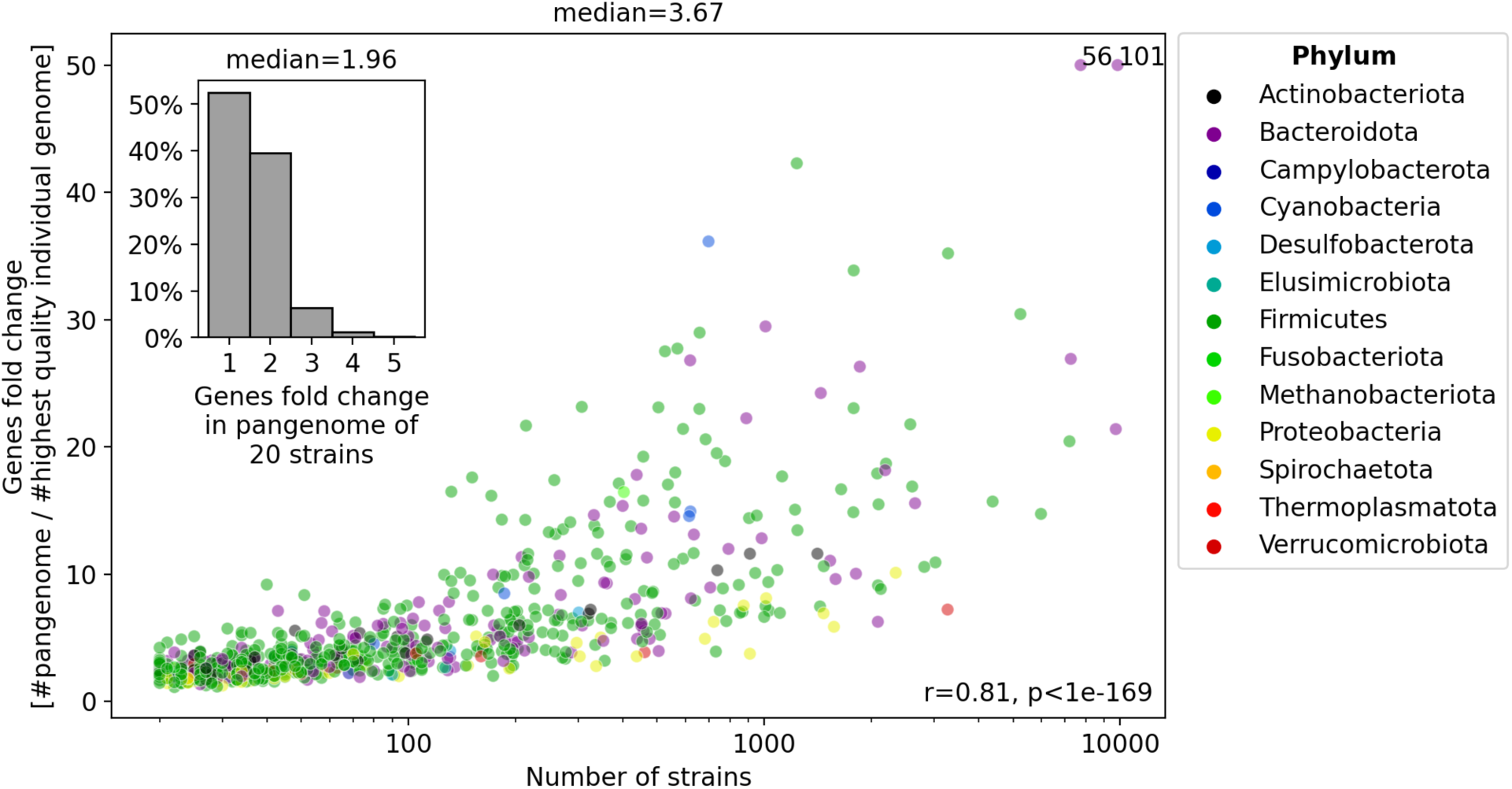
Pangenomes typically hold four times more genes than the highest quality individual genome. The figure illustrates the relationship between pangenomes and individual genomes sizes, quantified by the number of genes across 728 prokaryotic species found in the human gut. Each dot represents a species, with colors indicating their respective phyla (legend). The y-axis is the fold change in the number of genes within the pangenome compared to the highest quality individual genome of that species, while the x-axis shows the number of strains used to construct the pangenome (log10 scale). The median genes fold change is 3.67. For visualization purposes, two species with fold changes exceeding 50 are clipped, with their actual values provided next to them. Notably, a strong correlation exists between the fold change and the number of strains used to construct the pangenome (Spearman r=0.81, p<1e-169). To account for variation in strain availability, a histogram depicting the genes fold change obtained if only 20 strains were utilized to create each pangenome is shown (methods). In this scenario, the median genes fold change is 1.96.

### A balance between conserved and variable genes

We sought to investigate the extent of gene conservation within and between species. To this end, we categorized genes into three groups based on their frequency among strains of the same species; the core genes are defined as those present in 290% of strains, the shell genes are present in between 10%2 and <90%, and the cloud genes are present in <10% of strains (Figure 2).

**Figure 2.**
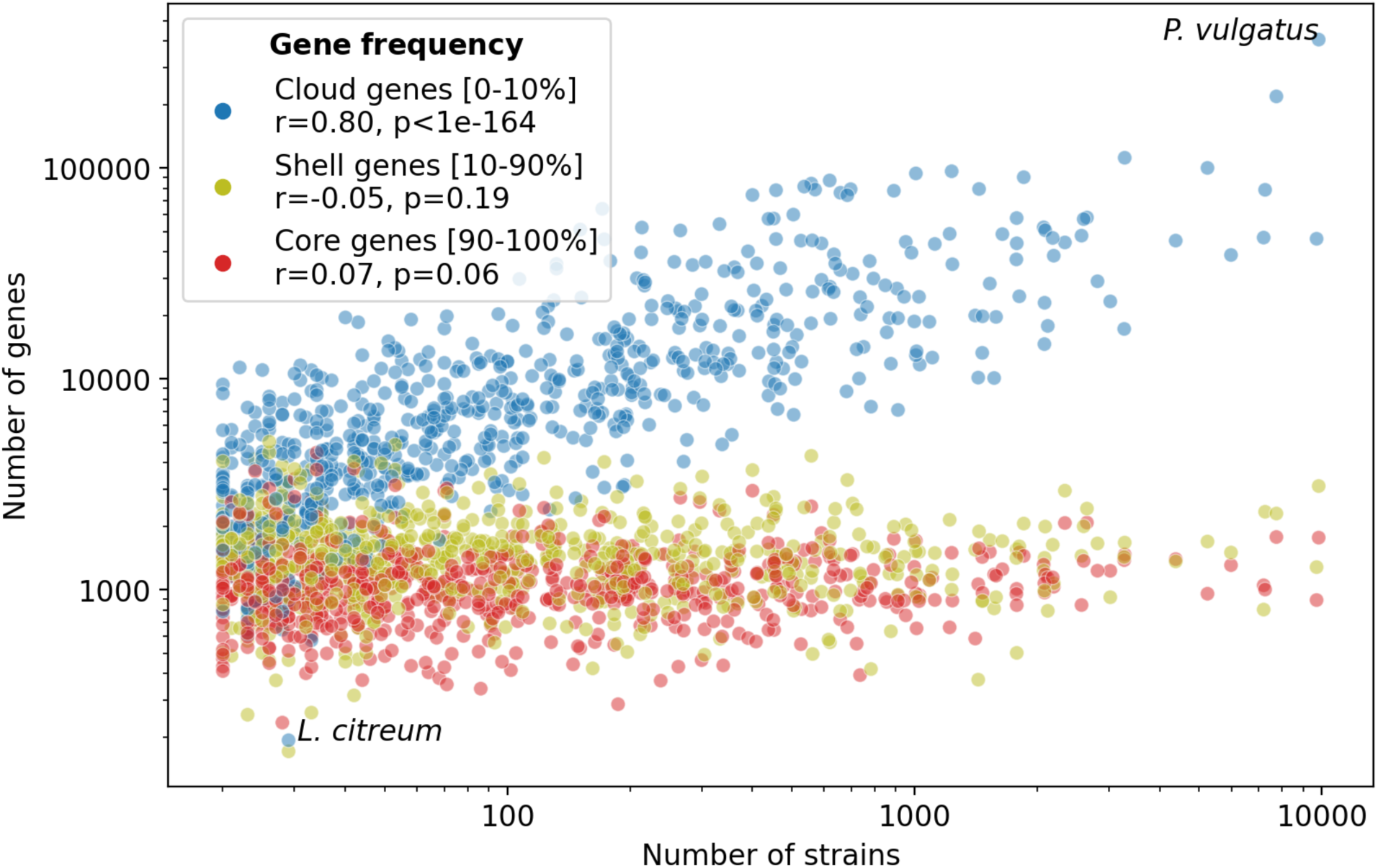
Species have a core set of a thousand genes that defines them, and an accessory set of genes that are unique to the different strains. The figure illustrates the gene composition within pangenomes of 728 prokaryotic species found in the human gut. The y-axis represents the number of genes in each gene frequency group (log10 scale), while the x-axis shows the number of strains used to construct the pangenome (log10 scale). Genes are categorized into three groups - core, shell, and cloud genes, based on their frequency across strains. The legend provides the frequency range and color of each group, with Spearman correlation between the number of strains and the number of genes in the group. Notably, the identity of core genes remains consistent regardless of the number of strains used to construct the pangenome (20 strains versus all available strains, Jaccard index 0.82±0.10, Figure S1, methods), while the number of cloud genes increases with added strains (Spearman r=0.80, p<1e-164). *Leuconostoc citreum* and *Phocaeicola vulgatus*, the species with the smallest and largest number of cloud genes, are highlighted, respectively.

We find that the typical species harbors 1,044 core genes (median, range 234-4,465), with no dependency on the number of strains used to construct the pangenome (Spearman r=0.07, p=0.06). Even when constructed from a limited set of 20 strains, pangenomes maintain a substantial overlap with the full pangenomes core genes (Jaccard index 0.82 mean ± 0.10 std, Figure S1, methods), underscoring the stability of core genes across varying sample sizes. However, when comparing species within the same genus, the core genes’ overlap diminishes markedly (Jaccard index 0.54±0.11, T-test for independent samples, p<1e-186), highlighting the distinctiveness even between closely related species. Remarkably, species belonging to the Bacteroidota phylum exhibit the highest degree of gene conservation compared to other major phyla in our dataset, Firmicutes, Actinobacteriota, and Proteobacteria (220 species, Jaccard index 0.36±0.07 versus 0.26±0.07, 0.25±0.10, and 0.25±0.11, respectively, Bonferroni corrected p<1e-224, Figure S1).

Similarly, the typical species holds 1,434 shell genes (median, range 171-5,025), with no dependency on the number of strains used to construct the pangenome (Spearman r=-0.05, p=0.19). This lack of dependency stems from the wide range of frequencies acceptable within this gene group - 10-90%, that vary with sample size (median Spearman r=-0.18±0.07, median p<1e-4 between the shell genes frequency in pangenomes constructed from 20 strains and in the full pangenomes, methods).

Conversely, the typical species has 6,649 cloud genes (median, range 193-111,694, with two outliers of 218,275 and 405,918), with strong dependency on the number of strains used to construct the pangenome (Spearman r=0.80, p<1e-164). Indicating considerable strain-level genetic diversity within the human gut microbiota.

Attempts to characterize the functional roles of each gene group were hindered by the low annotation rate resulting in untrustworthy results, especially in the cloud genes compared to shell and core genes (unannotated cloud genes 55.06%±7.70, shell 44.30%±9.94, and core 28.26%±5.28, T-test for independent samples, p<1e-215, Figure S2, methods).

One species, *Leuconostoc citreum*, stands out with only 193 cloud and 171 shell genes, but a reasonable number of 1,515 core genes. It also has the second smallest genes fold change, standing at 1.16. Looking into this species we found 27 out of the 29 strains came from the same neonatal study on days 4-16 of the infants lives ^7^. This may reflect a homogeneous source of the strains analyzed, and a limited presence in the general population (the other two strains came from elderly males). Indeed this species was detected in only seven out of 4,624 samples (0.15%) in validation cohorts from Israel and the Netherlands (in comparison the species with the highest number of cloud genes, *Phocaeicola vulgatus*, was detected in 4,457 of the samples - 96.39%) ^6^.

### Different species have different levels of strain variability

We wondered whether cloud genes discovery reached saturation in our dataset or if we would have more strains, we would keep discovering new cloud genes. To assess this, we quantified strain variability as the exponent in Heap’s law (“a”, methods) ^8^, which signifies whether gene discovery has reached a plateau (a=0, minimal gene discovery) or it continues to grow as new strains are introduced (a=1, maximal gene discovery). Our findings revealed considerable variation among species. While some species exhibit a saturated or “closed” pangenome, others maintain an “open” pangenome, indicating ongoing gene discovery (Figure 3). Strain variability has moderate correlation with species prevalence in the population (Spearman r=0.53, p<1e-53) and low correlation with species abundance among the individuals who have it in validation cohorts from Israel and the Netherlands (Spearman r=0.11, p=0.004) ^6^.

**Figure 3.**
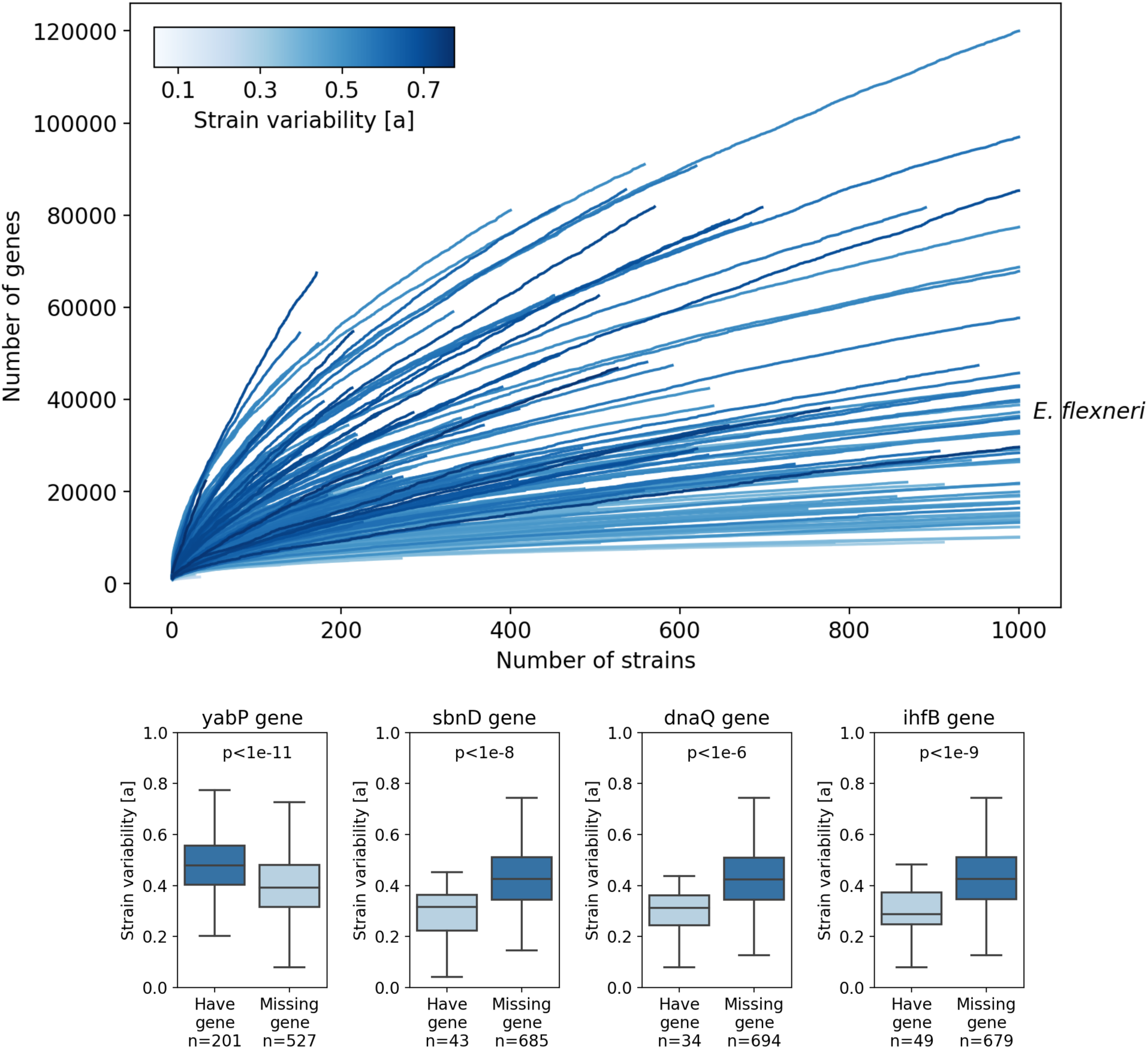
High strain variability associates with sporulation, whereas low strain variability associates with antibiotic resistance. The figure illustrates the different levels of strain variability across 728 prokaryotic species (different lines). The x-axis represents the number of strains (cut at 1,000 for visualization), while the y-axis shows the number of genes in the pangenome. The color indicates the level of strain variability, defined as the exponential dependency of gene count on the number of strains (methods). *Escherichia flexneri*, a close relative of *Escherichia coli*, is highlighted for reference. Boxplots show examples of genes associated with strain variability from the categories of sporulation (yabP), antibiotics resistance (sbnD), DNA repair (dnaQ), and DNA recombination (ihfB) (T-test for independent samples, Bonferroni corrected p-values shown, methods). Boxes show the quartiles of the data (0.25, median, 0.75), while the whiskers extend to 1.5 of the inter quartile range, points beyond the whiskers are considered to be outliers and are not shown.

**Heaps Law**: *N = k*n^a^*

Where N is the number of genes, n the number of strains, k constant of proportionality, and “a” exponent calculated to fit the formula.

To understand the mechanisms contributing to this variability, we analyzed the association between core genes and the exponent in Heap’s law. We find that out of 2,460 core genes tested, 191 showed significant association with strain variability (T-test for independent samples, Bonferroni corrected p<0.05, methods). Surprisingly, 34 of the 71 genes (48%) associated with high strain variability, and 7 out of the top 10, are related to sporulation (sporulation related genes ordered by significance - yabP, yabG, cwlD, spoI, sigE, ftsX, spoV, srrA, nrdG, mbl, sigG, whiA, porD, prkC, mgtE, yycJ, cwlJ, sigF, sigK, asnO, lytB, yerB, ydcC, lrgA, ytrA, divI, ctpB, ylmC, ywaC, cdsA, spmB, sspC, gdpP, ftsL) ^9-25^. We also identified four sporulation-related genes among the 120 (3%) associated with low strain variability (kdsB, anmK, yicL, dksA) ^9^.

Nine genes tied with low strain variability facilitate resistance to antibiotics (sbnD, lptD, menH, emrB, galE), organic solvents (mlaD, mlaC, mlaE), and Tellurite (htpX). While both sporulation and these resistance mechanisms protect bacteria, sporulation appears prevalent in species with high strain variability, while the resistance genes are found in those with low variability.

More expectedly, we observed seven genes linked with low strain variability involved in DNA repair (dnaQ, xthA, uvrD, nrdF, nrdE, rph, hfq), and two in DNA recombination (ihfB, ruvC). Another DNA recombination gene (recU) links to high strain variability.

To address potential confounding factors between the associated genes and phylogenetics, we compared the strain variability among the major phyla in our dataset (220 species). We found that Proteobacteria species, known to be highly antibiotic resistant ^26^, exhibit significantly lower strain variability compared to Firmicutes, Bacteroidota, and Actinobacteriota, underlining a distinctive aspect of its genomic diversity (0.29±0.08 versus 0.44±0.13, 0.43±0.09, and 0.39±0.06, respectively, T-test for independent samples, Bonferroni corrected p<1e-5, Figure S3, methods).

### Antibiotic resistance in the human gut

After identifying associations of low strain variability with Proteobacteria species and antibiotic resistance genes, we proceeded to map the landscape of antibiotic resistances within the human gut microbiota. We found 6,463 instances of 214 genes conferring resistance to 29 antibiotic classes within our pangenomes. There are 37 additional antibiotic drug classes in the database we searched against that we do not see at all (Figure 4, methods). Of the 728 species analyzed, 636 harbored at least one antibiotic resistance gene. These genes were mostly cloud genes (90.98% of the cases), indicating their sporadic presence across strains. However, in 237 drug-species pairs, resistance to the same antibiotic is carried by >90% of a species’ strains, from hereafter referred to as “extreme resistance”.

**Figure 4.**
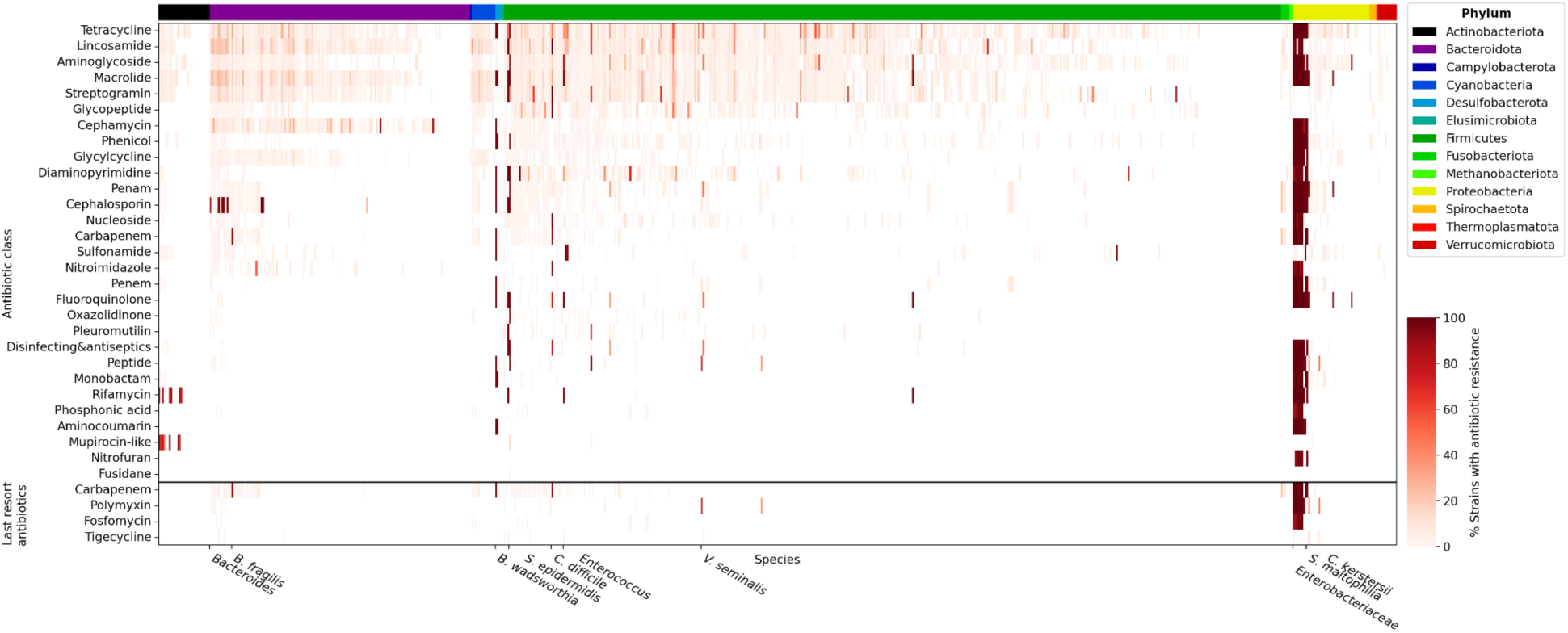
237 cases of extreme antibiotic resistance, predominantly of Enterobacteriaceae species. The heatmap illustrates the percentage of strains (right colorbar) of 728 human gut species (x-axis) harboring antibiotic resistance genes against specific antibiotic classes (y-axis). Species are organized alphabetically by their phylum (top colorbar and legend), and within each phylum, they are further sorted based on the number of antibiotic resistances detected. The antibiotic classes are sorted based on the number of species exhibiting resistance to them. The bottom panel highlights last resort antibiotics. Cephalosporins of the 4th and 5th generations, considered last resort antibiotics, are not included due to our limitation in distinguishing between Cephalosporin generations. Notably, extreme antibiotic resistances are predominantly observed within species of the Enterobacteriaceae family, even to last resort antibiotics, underscoring the critical health threat posed by these bacteria.

Among these cases, 174 (73%) belong to Proteobacteria species, with 136 (57%) within the highly antibiotic-resistant Enterobacteriaceae family, 16 of *Stenotrophomonas maltophilia*, and 14 of *Comamonas kerstersii* (another three species have two to three extremes resistances). Remarkably, some of the Enterobacteriaceae species exhibited extreme resistance to as many as 22 antibiotic classes, including the last resort antibiotic Carbapenem, posing a major health threat ^26-28^. The six antibiotic classes we do not see Enterobacteriaceae resistance against (Fusidane, Glycopeptide, Mupirocin-like, Oxazolidinone, Pleuromutilin, Streptogramin) target Gram-positive bacteria while Enterobacteriaceae species are Gram-negative, and thus are not susceptible and have not developed resistance to these drugs ^29-30^. The low resistance of Enterobacteriaceae species to Sulfonamide, that does target Gram-negative bacteria, could potentially be explained by trends in prescribing practices where newer antibiotics are favored over older ones like Sulfonamide ^31^, and consequently species that are less exposed to Sulfonamide also carry less resistance to it. Interestingly, even though the two species outside of the Enterobacteriaceae family have similar amount of resistances, *S. maltophilia* is known to cause numerous infections ^32-33^, while reports of infections due to *C. kerstersii* have been rare in the past decade ^34-39^. This phenomenon might be explained by related species being susceptible and subsequently developing resistances to the same drugs, even if those drugs are meant to target other closely related species.

Following Proteobacteria, are Desulfobacterota species *Bilophila wadsworthia* and Firmicutes species *Staphylococcus epidermidis* exhibiting 14 and 11 extreme resistances, respectively. *B. wadsworthia* is frequently isolated from hospital-related infections, particularly post-surgery, with varying antibiotic susceptibility ranges across different geographical regions - for example selective antibiotic susceptibility in the Netherlands and almost no susceptibility in Japan ^40-41^. *S. epidermidis*, often associated with medical devices, exhibits variable antibiotic susceptibility, which significantly drops in hospital environments ^42^, in line with our results.

From an antibiotic standpoint, 19 species have extreme resistance to Cephalosporin, 17 species to Macrolide, and 17 species to Fluoroquinolone (ten, six, and five of the species are not Proteobacteria, respectively). Most of the non-Proteobacteria extreme resistances to Cephalosporin are of species from the Bacteroides genus, and the resistances to Macrolide and Fluoroquinolone from the Enterococcus genus. All three antibiotic resistances are confirmed by the literature ^43-45^.

Particularly concerning is the emerging resistance to last resort antibiotics, kept for cases where traditional treatments have failed ^46^. We find seven Enterobacteriaceae species with extreme resistance to the last resort antibiotic Polymyxin, six to Carbapenem, and four to Fosfomycin, with two other species very close by - currently holding resistance to Fosfomycin in 89% of their strains. Fosfomycin is often used orally for urinary tract infections but is considered a last resort antibiotic when used intravenously against systemic infections ^47^. According to our data, Enterobacteriaceae species are most susceptible to the last resort antibiotic Tigecycline, approved by the United States food and drug administration (FDA) in June 2005 ^48^. Other than Enterobacteriaceae species, *S. maltophilia*, *C. kerstersii*, and *B. wadsworthia* also developed extreme resistance to Carbapenem, while *Bacteroides fragilis* (89%) and *Clostridioides difficile* (87%) are nearly as resistance. *Veillonella seminalis* is near extreme resistance (83%) to Polymyxin. Although Cephalosporins of the 4th and 5th generations are considered last resort antibiotics, our annotations did not provide sufficient specificity to distinguish between generations.

This analysis underscores the complex landscape of antibiotic resistances within the human gut microbiota. Considering most of our individual genomes came from generally healthy non-hospitalized individuals, further highlights the pressing need for continued surveillance and development of new treatments to combat this emerging threat.

### Strain association with host characteristics

With increasing evidence linking microbial species composition to host characteristics, we explored whether strain specific genes are involved in this connection. For 200 species with 2200 strains, we clustered the strains to delineate species sub-groups based on the presence-absence information of the shell genes (methods, Figure 5). We chose the shell genes because they exhibit the most variability within a species, while the core genes are present and the cloud genes are absent in most strains. In cases where sub-species groups were found, we investigated whether they associate with the age or sex of the host. We found 24 species in which the sub-groups significantly associate with age, and five with the sex of the human host (linear model, Bonferroni corrected p<0.05, Table S1, methods). In three species - *Akkermansia muciniphila*, *Phocaeicola vulgatus*, and a species of the Acutalibacteraceae family - the sub-species groups showed significant associations with both age and sex.

**Figure 5.**
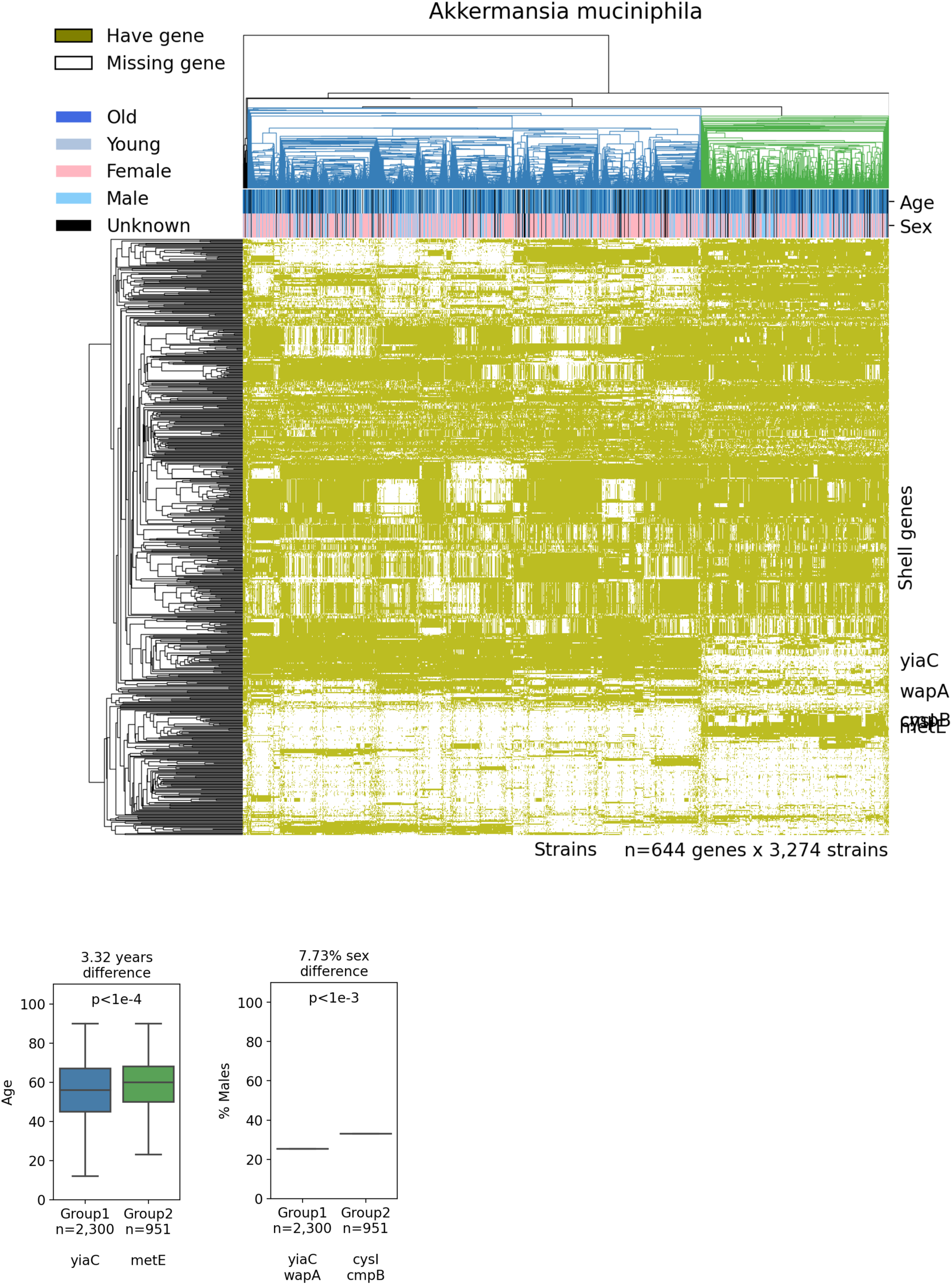
*Akkermansia muciniphila* strains associate with host age and sex. The clustermap illustrates sub-species groups (color in the upper dendrogram) based on the presence (olive color) and absence (white color) of shell genes (y-axis) in the strains (x-axis) (showing only sub-group significantly associated genes according to a linear model, Bonferroni corrected p<0.05, methods). Host age and sex are indicated in the upper colorbars and in the boxplots (linear model, Bonferroni corrected p shown, methods). Genes associated with a sub-group and a phenotype are listed below the corresponding box. Boxes show the quartiles of the data (0.25, median, 0.75), while the whiskers extend to 1.5 of the inter quartile range, points beyond the whiskers are considered to be outliers and are not shown.

People carrying sub-species group1 of *A. muciniphila* are 3.32 years younger and 7.73% more likely to be females than people carrying sub-species group2 (p<1e-4 and p<1e-3, respectively, Figure 5). We found numerous supporting evidence for the connection between *A. muciniphila* abundance and the age ^49-54^ and sex ^54-59^ of the host, and sought to understand whether these associations are affected by the strain-level genetic differences. To this end, we tested which of the shell genes associated with the sub-groups are also associated with age or sex. We found that out of 644 shell genes associated with the *A. muciniphila* sub-groups, 35 also associate with the age of the host (linear model, Bonferroni corrected p<0.05, methods). Only two of these genes are annotated - the yiaC and metE genes. yiaC is a lysine acetyltransferase ^60^. The acetylation of lysine side chains is a post-translational modification, conserved from bacteria to human, that regulates fundamental cellular processes with known implications on the organisms’ aging process: metabolism, transcription, translation, cell proliferation, regulation of the cytoskeleton, and DNA repair ^61^. Recently identified enzymes with dual deubiquitinase and acetyltransferase activity injected by bacteria into host cells showed that post-translational lysine acetylation is used by the bacteria to modulate host cell processes ^61^. In our analysis, people carrying *A. muciniphila* with the yiaC gene are younger than those without the gene. The other gene, metE, catalyzes the transfer of a methyl group from 5-methyltetrahydrofolate to homocysteine resulting in methionine formation. Methionine-rich diets, possibly mediated by the microbiota, have demonstrated accelerated aging-related changes, including increased vascular oxidative stress, mitochondrial dysfunction, inflammation, and impaired cognitive performance ^62^. In our analysis, people carrying *A. muciniphila* with the metE gene are older than those without the gene.

We additionally found 40 *A. muciniphila* shell genes that associate with the sex of the host (linear model, Bonferroni corrected p<0.05, methods). Only four of these genes are annotated - the yiaC (as before), wapA, cysI and cmpB genes. In regards to yiaC, lysine acetylation of androgens (a set of sex hormones produced in both males and females) receptors plays a vital role in directing its cellular activities via modulating its stability, nuclear localization, and transcriptional activity ^63^. In our analysis, people carrying *A. muciniphila* with the yiaC gene are more likely to be females than those without the gene. cysI is a nitrite and sulphite reductase. Females have higher nitrite levels and greater nitrate reduction capability, found to be modulated by the oral microbiota ^64^. Sulfite reductase is the reduction of sulfite to sulfide, a product that is often protonated to form hydrogen sulfide. Hydrogen sulfide affects sexual function in both males (pro erectile relaxant effect on the cavernosum) and females (smooth muscle relaxant effect in genital tract) ^65-66^. In our analysis, people carrying *A. muciniphila* with the cysI gene are more likely to be males than those without the gene. However, in both subgroups, the chance of being a female is higher than the chances of being a male. We did not find literature to support the connection between wapA and cmpB genes to the sex of the host.

Similarly, people carrying sub-species group1 of *P. vulgatus* are 1.67 years younger and 8.19% more likely to be females than people carrying sub-species group2 (p<1e-3 and p<1e-12, respectively, Figure S4). We found support connecting *P. vulgatus* abundance and the age ^67-68^ and sex ^69^ of the host. Out of 2,596 shell genes associated with the *P. vulgatus* sub-groups, 350 also associate with the age of the host (linear model, Bonferroni corrected p<0.05, methods). 85 of these genes are annotated, and of them 30 (35.29%), including 24 out of the top 30, remarkably encode for ribosomal proteins (18 large and 12 small ribosomal subunit proteins).

Reducing ribosomal proteins heterogeneity negatively affects the rate and precision of protein biosynthesis a phenomena linked with older age in many organisms ^70-73^. Two other protein synthesis related genes came up - infA and tufA, in addition to two protein folding genes - surA and groL. In our analysis, people carrying *P. vulgatus* with most of the ribosomal genes are younger than those without the genes. It is possible that the loss of ribosomal proteins with increasing age is related to the cumulative use of antibiotics over a person’s lifespan. Some antibiotics target ribosomes or proteins involved in their assembly and function, and in cases where they are not essential for bacterial survival, it could have lost ribosomal proteins in order to not be susceptible to the drug. We additionally found 775 genes that associate with the sex of the host (linear model, Bonferroni corrected p<0.05, methods). 146 of the genes are annotated. The top annotated genes are isiB, dapb3 and pehX genes, however we did not find literature supporting the connections between them and sex. Overall, these findings highlight the intricate relationship between microbial genetics and the human host.

## Discussion

In this work, we created pangenomes for 728 human gut prokaryotic species, the majority of which are bacteria, revealing a vast genetic diversity that underscores the adaptability and resilience of the biota. Our findings, encompassing the significant expansion in gene count within pangenomes compared to individual genomes, the balance between conserved and variable genes, and the distinct functional roles and diversity across species, provide crucial insights into microbial evolution and the potential impact on host health.

The discovery of a four fold increase in gene content within pangenomes compared to individual species genomes underscores the remarkable genetic capacity of the gut microbiota. This expansion is indicative of the rich functional diversity that gut prokaryotes can harbor, contributing to various metabolic pathways and interactions within the gut ecosystem. The stability of core genes across varying numbers of strains, alongside the marked distinction in gene conservation between species within the same genus, indicates a complex interplay of evolutionary forces shaping the genetic diversity of the gut microbiota. It is possible the high core genes conservation seen within the Bacteroidota phyla and low strain variability seen within the Proteobacteria phyla, reflect their later divergence in the tree of life compared to the early divergence of Firmicutes and Actinobacteriota phylum ^74^.

The association between specific functions and strain variability, particularly the link of high variability with sporulation, suggests an evolutionary strategy of temporal and spatial “skipping” for survival. Sporulation allows bacteria to withstand a broad range of unfavorable conditions, which may reduce the evolutionary pressure they need to withstand, resulting in strain variability maintenance. The presence of antibiotic resistance genes, associated with low strain variability, highlights a recent adaptation of the bacteria to a specific type of pressure, leaving them susceptible to other unfavorable conditions that may drive their strain variability loss.

Other studies support the associations found with these specific functions. One study showed that upon niche migration, an event enabled by sporulation, genetic diversity introduced by horizontal gene transfer can be maintained, and is otherwise lost ^75^. Another study supported this notion by showing spore formation is required for gut bacteria to transmit between non-cohabiting humans ^76^. A third study suggested that because the human microbiota is globally connected, local contamination of the mobile gene pool by antibiotic resistance genes can have significant transnational consequences ^5^.

The mapped landscape of antibiotic resistance genes within the human gut microbiota, of mostly healthy individuals, shows the majority of species carry resistance to at least one class of antibiotics. These genes are usually cloud genes, indicating their sporadic presence across strains. However, in Proteobacteria and several other species extreme resistance (carried by >90% of strains) to multiple drugs is seen, including to last resort antibiotics like Carbapenem. These species pose a significant public health risk, underscoring the urgency for surveillance and the development of novel treatment strategies. One such strategy could be to discontinue the use of certain antibiotics and then reinstate them once the resistance in the population diers, like we see in the case of Sulfonamide, which has been less prescribed in recent years ^31^.

The analysis of strain association with host characteristics further illuminates the intricate interactions between gut microbiota and human health. The significant associations between specific strains and host age and sex suggests unchangeable host characteristics can influence microbial community composition at the gene level. A theoretical example of such influence could be mediated by androgens. Androgens, typically present in much higher levels in males than in females, and in young females more than in post-menopausal females, can inflict evolutionary pressure on the bacteria that strengthen strains that carry genes that modulate the activity of androgen receptors in the bacteria more efficiently, in line with our results. The findings regarding *Akkermansia muciniphila* and *Phocaeicola vulgatus* for example, offer promising avenues for personalized microbiota-based interventions, potentially targeting age-related health issues and sex-specific health outcomes. For example, attenuating methionine producing or ribosomal heterogeneity lacking strains, might be able to reduce the metabolic decline associated with aging.

Our study faces several limitations, including the reliance on metagenomic derived genomes that are often less complete and more contaminated than isolates derived genomes. However, our genomes are estimated to be 91% complete and only 1% contaminated (medians) ^6^, and enable the study of uncultivated species, and obtain a large set of genomes. With that said, we are still limited by the available genomic data, which may not fully capture the diversity of the human gut microbiota across different populations. Additionally, the functional roles of many genes, especially within the cloud gene category, remain poorly understood due to low annotation rates. This gap in knowledge highlights the need for further research to elucidate the functional implications of microbial genetic diversity for human health.

In conclusion, our work provides a foundational understanding of the genetic diversity within the human gut microbiota and its implications for human health. The balance between conserved and variable genes reflects the complex evolutionary history of gut prokaryotes and their adaptation to the gut environment. The prevalence of antibiotic resistance genes emphasizes the need for vigilant monitoring and innovative treatment strategies. Finally, the association between microbial genes and host characteristics opens new avenues for personalized medicine, leveraging the gut microbiota for health optimization. Further research is needed to translate these insights into practical applications, ultimately improving human health through targeted microbiota interventions.

## Methods

### Genome assemblies

Genome assemblies of human gut prokaryotes were sourced from the Weizmann Institute of Science (WIS) genome set ^6^. This set encompasses 241,118 genomes meeting the quality criteria of completeness 270% and contamination :55% ^77^. In cases where multiple genomes from the same individual were highly similar (:50.05 genomic distance apart), only the highest quality genome was retained (based on a quality score described below).

To organize these genomes into meaningful biological groups, species, genus, and family clusters were set based on average genomic similarity thresholds of 0.05, 0.15, and 0.30 MiniHash (MASH) distance, respectively ^78-79^. MASH distance is a good proxy for one minus the average nucleotide identity (ANI), so that a MASH distance of 0.05 corresponds to the widely recognized prokaryotic species delineation standard of 95% ANI ^2^.

Within each species cluster, the best genome was identified using a composite quality score, which balances completeness, contamination, and N50 contig size as follows ^6^: *quality score = completeness - 5*contamination + 15*log_10_(N50)* Where N50 is the contig size at which half of the total genome length is covered by contigs of equal or greater size. Taxonomy was assigned to the best species’ genomes using the Genome Taxonomy Database (GTDB) toolkit ^80^.

Of the 3,594 species in the WIS genome set, 728 species, comprising 725 bacterial and three archaeal species, possessed 220 genomes within their species-level clusters and thus were included in our analysis.

### Pangenomes construction

Pangenomes, representing the collective gene pool of different strains within a species, were constructed based on gene clustering, conserved gene neighborhood information, and weighted graphs, using a common program called Roary ^81^ ^8^. Roary version 3.13.0, with the following arguments: -p 32 number of threads, -e create a multiFASTA alignment of core genes using PRANK, -n fast core gene alignment with MAFFT, -i 90 minimum percentage identity for blastp, -cd 90 percentage of isolates a gene must be in to be core, -g 1000000 maximum number of clusters, -s do not split paralogs. Roary’s input files in GFF format were created from the WIS genome fastas ^6^ using Prokka ^82^ version 1.14.6, with the following arguments: --cpus 2 number of CPUs to use, --rfam enable searching for ncRNAs with Infernal+Rfam.

The number of genes in a pangenome constructed from a subset of genomes is dependent on the set of genomes chosen. To mitigate this selection bias, whenever a subset was used we averaged the gene count from ten randomly chosen subsets of genomes.

### Gene annotations

Gene annotations were produced using Prokka as mentioned above, and eggNOG ^83^ version 2.0.4, with the following arguments: --dbmem option to pre-load the eggnog.db sqlite3 DB into memory, --go_evidence all option to report all GO terms, -m diamond search queries against eggNOG sequences using diamond, --cpu 16 number of CPUs to be used whenever possible, --itype CDS the type of sequences included in the input file. Gene’s functional categories were extracted from the eggnog output.

Antibiotic resistance annotations were produced using ABRicate ^84^ version 1.0.1, with the following arguments: -db CARD ^85^ (pulled on February 8th, 2024) database to use, -minid 70 minimum DNA %identity, -mincov 70 minimum DNA %coverage. A strain is said to be resistant to an antibiotic if it has any number of genes conferring resistance to it.

For the largest species in our work (ID 449), Roary generated gene presence/absence information but failed to create a pangenome fasta file, precluding this species from eggNOG and ABRicate based analyses.

### Strain variability

Pangenome openness was quantified in the common practice by the exponent in Heaps Law ^8^:

N = k*n^a^

Where N is the number of genes, n the number of genomes, k constant of proportionality, and “a” exponent calculated to fit the formula.

Associations between functions and pangenome openness were assessed using annotated core genes, no matter the copy variation, present in 230 genomes and absent from 230 genomes. The reasons for only considering core genes are 1) they are characteristic to most genomes of the species, and 2) they have similar amounts of genes in both “open” and “closed” pangenomes, thus not biasing the results as “open” pangenomes have higher number of genes.

### Strain association with host characteristics

Sub-species were delineated within species with 2200 genomes based on gene presence/absence patterns, using average Euclidean distance for genes, and average MASH distance for genomes ^79^. Sub-species were delineated as the primary group of genomes at the apex of the clustering dendrogram, comprising 250 entities but :580% of all the species entities, displaying a genetic divergence of at least 1% from their parent branch.

Our focus on shell genes stemmed from their dynamic nature compared to core and cloud genes within species. Core genes are ubiquitously present across most genomes, while cloud genes are absent from the majority of genomes. In contrast, shell genes exhibit great variance within species, offering insights into intra-species diversity. Sub-species association with shell genes was assessed using a Logit model. Sub-species association with human age and sex was assessed using Ordinary Least Squares (OLS) and Logit models, respectively. We had available age and sex information for 92% and 94% of our genomes, respectively.

### Statistical information

A two-sided Bonferroni corrected significance threshold of alpha=0.05 was applied to all statistical tests.

## Code availability

Computational analysis was performed in python version 3.7 using the following packages: numpy version 1.21.0 and pandas version 1.2.5 for data processing, scipy version 1.7.0 statsmodels version 0.13.2 and statannot version 0.2.3 for statistical analyses, matplotlib version 3.4.3 and seaborn version 0.12.0 for plotting.

## Supplementary

**Figure S1.**
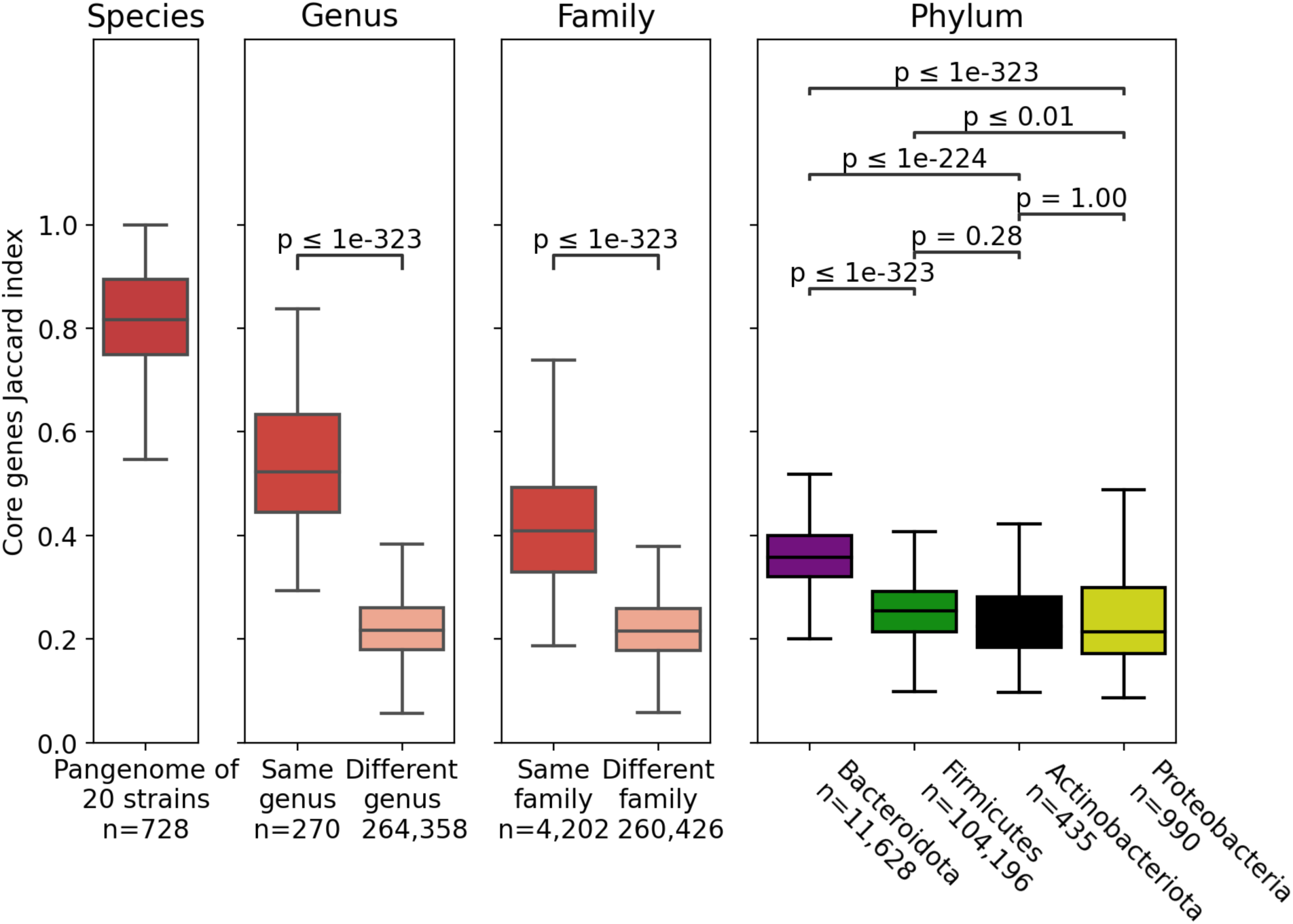
Core genes are conserved even in small sample size, and diminish markedly within closely related species. **a.** Core genes conservation between the full pangenomes and those constructed from only 20 strains (methods). **b, c, and d.** Core genes conservation within species from the same and from different genera, families, and phylum, respectively (for phyla with 220 species, sorted by the median value, species comparison to self are excluded, T-test for independent samples, Bonferroni corrected p-values shown in d, methods). Boxes show the quartiles of the data (0.25, median, 0.75), while the whiskers extend to 1.5 of the inter quartile range, points beyond the whiskers are considered to be outliers and are not shown. Notably, Bacteroidota species exhibit the highest degree of core genes conservation compared to the other big phyla.

**Figure S2.**
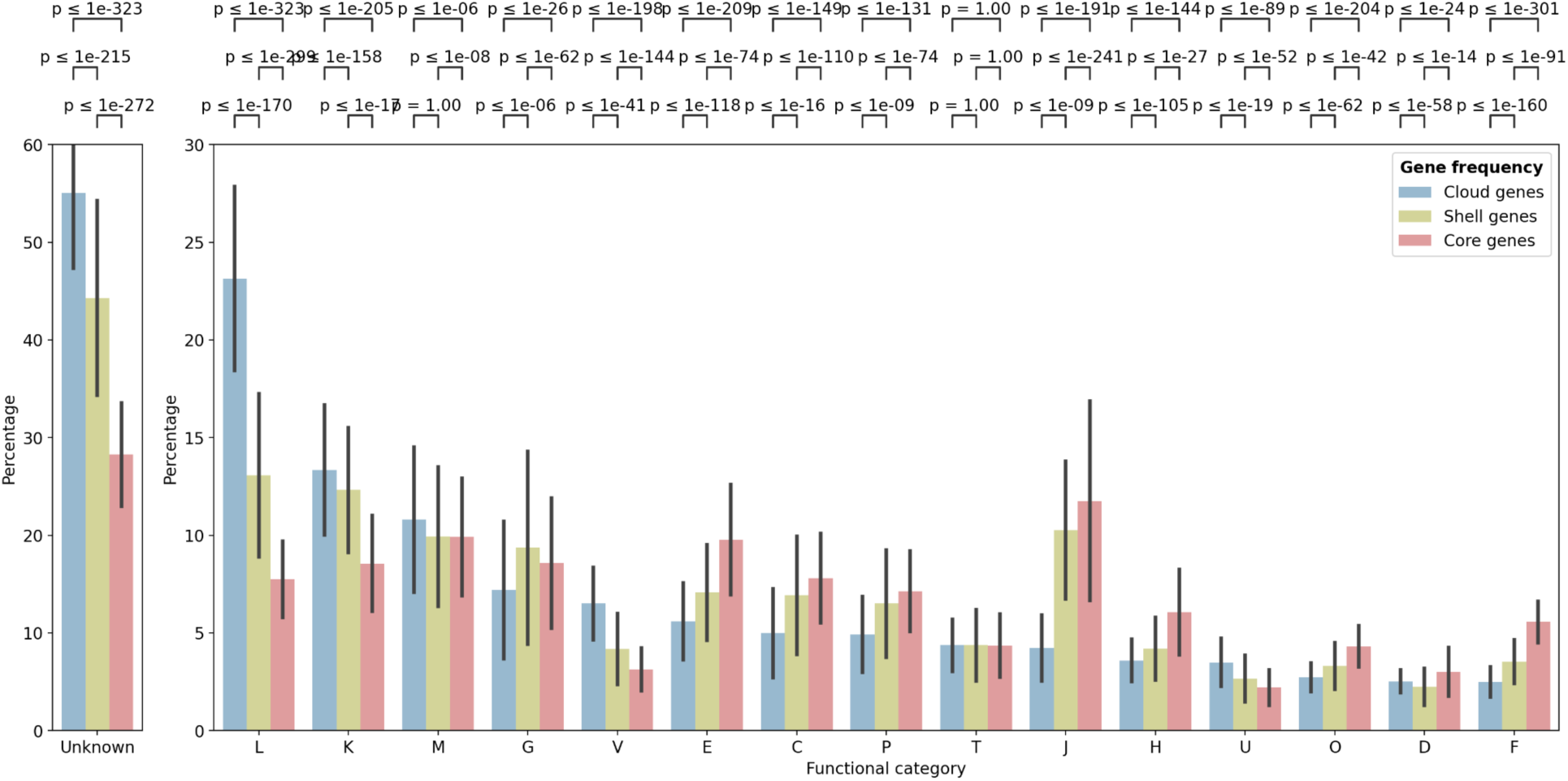
Functional categories across gene frequency groups. For each gene frequency group (legend), the y-axis denotes the percent of genes with unknown function (left panel), and with a known function (right panel). The x-axis lists functional categories (detailed below) that constitute 21% of the annotations (T-test for paired samples, Bonferroni corrected p-values shown). The bar height indicates the mean across 727 species, and the error bar the standard deviation (excluding one species for which this kind of annotations could not be generated, methods). Notice the left and right panels’ y-axes are in different scales. For example, functions such as replication (L), a crucial biological process, shows a disproportionate representation among cloud genes compared to core genes. This unexpected finding leads us to question the reliability of this analysis, presumably caused by the significant proportion of unannotated genes. **L** - Replication and repair **K** - Transcription **M** - Cell wall/membrane/envelope biogenesis **G** - Carbohydrate metabolism and transport **V** - Defense mechanisms **E** - Amino acid metabolism and transport **C** - Energy production and conversion **P** - Inorganic ion transport and metabolism **T** - Signal transduction **J** - Translation **H** - Coenzyme metabolism **U** - Intracellular trafficking and secretion **0** - Post-translational modification, protein turnover, and chaperone **D** - Cell cycle control and mitosis **F** - Nucleotide metabolism and transport

**Figure S3.**
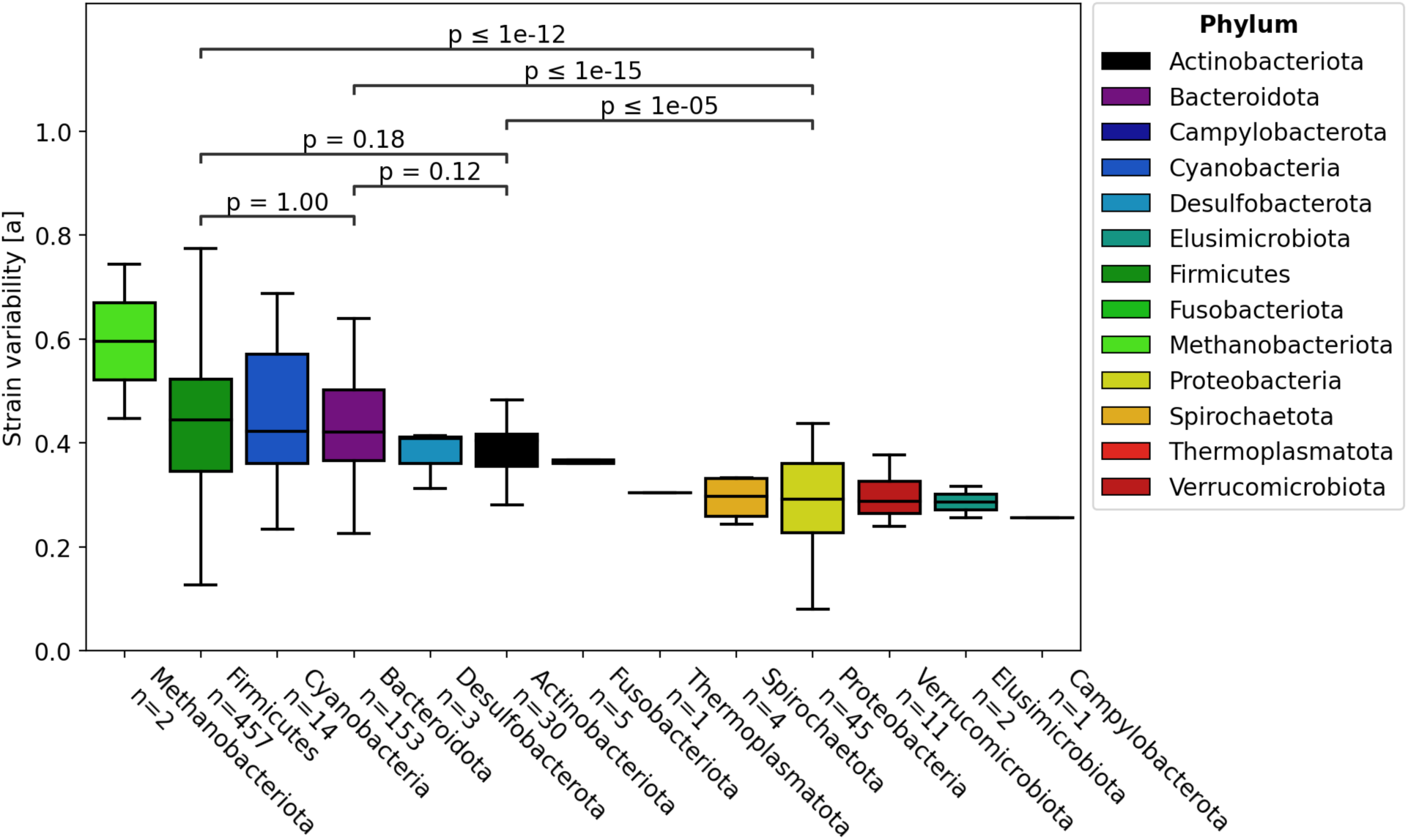
Proteobacteria species have less strain variability. The figure illustrates the strain variability (y-axis) across different phyla, sorted by median value (T-test for independent samples in phyla with 220 species, Bonferroni corrected p-values shown, methods). Boxes show the quartiles of the data (0.25, median, 0.75), while the whiskers extend to 1.5 of the inter quartile range, points beyond the whiskers are considered to be outliers and are not shown. Notably, Proteobacteria species exhibit less strain variability compared to other major phyla.

**Figure S4.**
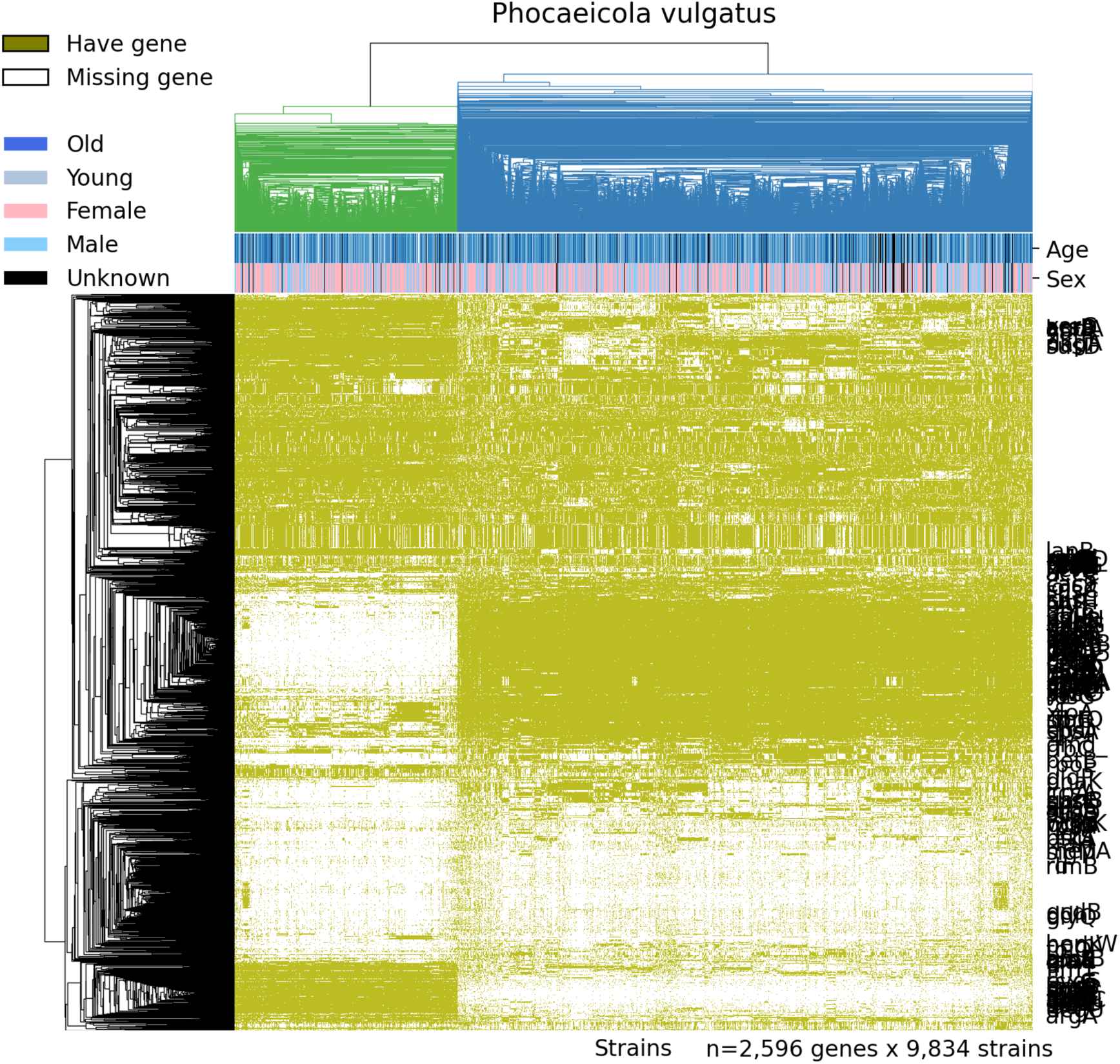
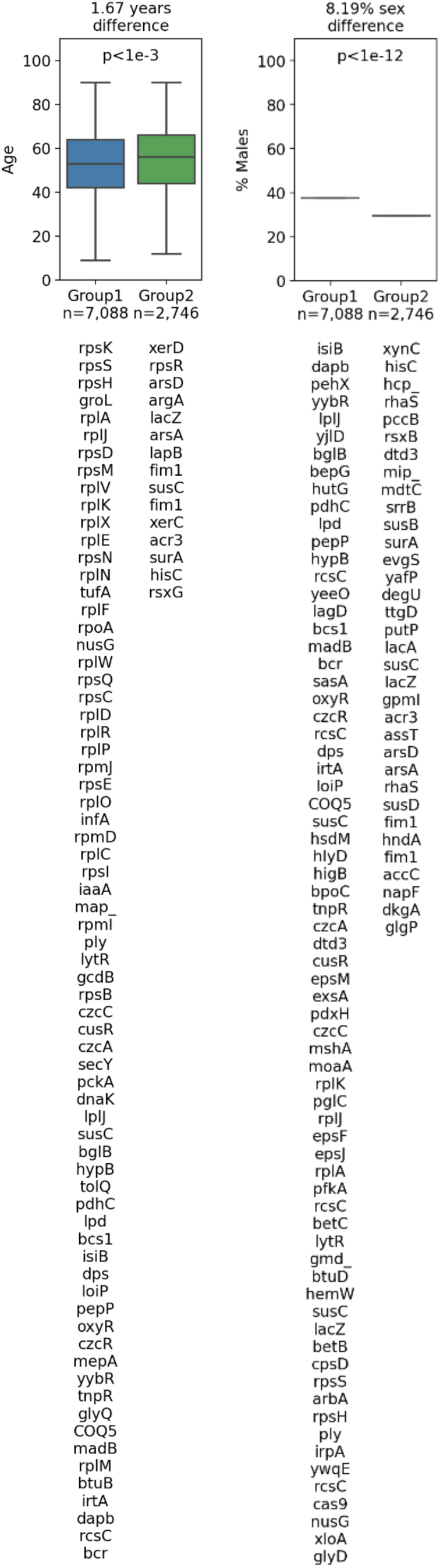
*Phocaeicola vulgatus* strains associate with host age and sex. The clustermap illustrates sub-species groups (color in the upper dendrogram) based on the presence (olive color) and absence (white color) of shell genes (y-axis) in the strains (x-axis) (showing only sub-group significantly associated genes according to a linear model, Bonferroni corrected p<0.05, methods). Host age and sex are indicated in the upper colorbars and in the boxplots (linear model, Bonferroni corrected p shown, methods). Genes associated with a sub-group and a phenotype are listed below the corresponding box, group1 sex genes are cut to fit the page. Boxes show the quartiles of the data (0.25, median, 0.75), while the whiskers extend to 1.5 of the inter quartile range, points beyond the whiskers are considered to be outliers and are not shown.

**Table S1.** Strain associations with host characteristics.

| **Species ID** | **Taxonomy** | **Phenotype** | **Effect size**<br>**[years or males]** | **Bonferroni**<br>**corrected p-value** |
| --- | --- | --- | --- | --- |
| 51 | *Bacteroidales* | Age | -10.47 | 5.00E-19 |
| 54 | *Bacteroidales* | Age | 14.12 | 6.00E-11 |
| 124 | *Prevotellamassilia* | Age | 18.9 | 1.00E-13 |
| 215 | *Prevotella* | Age | 9.43 | 3.00E-09 |
| 216 | *Prevotella stercorea* | Age | 9.46 | 3.00E-10 |
| 231 | *Prevotella copri* | Sex | -0.05 | 5.00E-05 |
| 280 | *Muribaculaceae* | Age | -10.03 | 2.00E-19 |
| 449 | *Phocaeicola vulgatus* | Age | -1.67 | 4.00E-06 |
| 449 | *Phocaeicola vulgatus* | Sex | 0.09 | 2.00E-15 |
| 620 | *Alistipes onderdonkii* | Age | -4.31 | 7.00E-05 |
| 721 | *Akkermansia muciniphila* | Age | -3.32 | 3.00E-07 |
| 721 | *Akkermansia muciniphila* | Sex | -0.09 | 1.00E-06 |
| 722 | *Akkermansia muciniphila B* | Age | -7.52 | 0.0002 |
| 959 | *Escherichia flexneri* | Age | 5.79 | 6.00E-07 |
| 1011 | *Succinivibrio* | Age | 7.04 | 7.00E-19 |
| 1211 | *Bacilli* | Age | -8.71 | 8.00E-17 |
| 1229 | *Catenibacterium* | Age | 17.57 | 7.00E-10 |
| 2185 | *Agathobacter rectalis* | Age | 2.92 | 1.00E-07 |
| 2216 | *Acetatifactor* | Age | 10.5 | 4.00E-07 |
| 2217 | *Acetatifactor* | Age | -20.93 | 3.00E-21 |
| 2378 | *Butyrivibrio A crossotus* | Sex | -0.17 | 0.0001 |
| 3056 | *Gemmiger formicilis* | Age | -10.47 | 0.0002 |
| 3078 | *Faecalibacterium prausnitzii* | Age | 9.13 | 8.00E-08 |
| 3216 | *Oscillospiraceae* | Age | -9.28 | 0.0001 |
| 3229 | *Oscillospiraceae* | Age | 4.82 | 5.00E-05 |
| 3523 | *Eubacterium R* | Age | 6.56 | 1.00E-06 |
| 3533 | *Acutalibacteraceae* | Age | 5.75 | 0.0001 |
| 3541 | *Acutalibacteraceae* | Age | 1.68 | 3.00E-06 |
| 3541 | *Acutalibacteraceae* | Sex | 0.2 | 4.00E-11 |

## Notes

### Competing Interest Statement

E.S. is a paid consultant to Pheno.AI, Ltd, other authors declare no competing interests.

